# A systematic screen for co-option of transposable elements across the fungal kingdom

**DOI:** 10.1101/2023.10.20.563377

**Authors:** Ursula Oggenfuss, Thomas Badet, Daniel Croll

## Abstract

How novel protein functions are acquired is a central question in molecular biology. Key paths to novelty include gene duplications, recombination or horizontal acquisition. Transposable elements (TEs) are increasingly recognized as a major source of novel domain-encoding sequences. However, the impact of TE coding sequences on the evolution of the proteome remains understudied. Here, we analyzed 1,237 genomes spanning the phylogenetic breadth of the fungal kingdom. We scanned proteomes for evidence of co-occurrence of TE-derived domains along with other conventional protein functional domains. We detected more than 13,000 predicted proteins containing potentially TE-derived domain, of which 825 were identified in more than five genomes, indicating that many host-TE fusions may have persisted over long evolutionary time scales. We used the phylogenetic context to identify the origin and retention of individual TE-derived domains. The most common TE-derived domains are helicases derived from *Academ*, *Kolobok* or *Helitron.* We found putative TE co-options at a higher rate in genomes of the Saccharomycotina, providing an unexpected source of protein novelty in these generally TE depleted genomes. We investigated in detail a candidate host-TE fusion with a heterochromatic transcriptional silencing function that may play a role in TE and gene regulation in ascomycetes. The affected gene underwent multiple full or partial losses within the phylum. Overall, our work establishes a kingdom-wide view of how domains likely derived from TEs contribute to the evolution of protein functions.

## Introduction

Proteomes are diverse and variability extends to the population and individual level [1]. Causes of proteome diversity include alternative splicing, presence-absence polymorphisms, single nucleotide polymorphisms or larger structural variations, such as duplications, reshuffling of protein domains, partial deletions or translocations [2]. Aneuploidy or gene duplication, followed by neofunctionalization due to relaxed purifying selection, can lead to diversification [3]. Gene gains can also be mediated by horizontal gene transfer from other species or by *de novo* gene birth [4,5]. Proteome evolution can also entail pseudogenization, with pseudogenes expected to eventually get lost or regain function. Genetic variation can provide a highly dynamic proteome, allowing populations to rapidly adapt to new or changing environments.

Mutations, rearrangements, losses or acquisitions of protein-coding genes may be facilitated by co-localization with transposable elements (TEs). In some species, TEs are clustered into gene-poor islands [6,7]. TE rich islands are under relaxed purifying selection, often leading to retention of single nucleotide polymorphisms or structural variants, and a higher rate of ectopic recombination caused by repetitive sequences [8]. Genes located in such TE islands are thought to be under purifying selection as well, allowing the accumulation of mutations at a fast rate [9]. Ectopic recombination of two copies of the same TE family can lead to the deletion of adjacent genes [10]. Some TEs such as *Helitrons* in maize or *Pack-Mules* in rice have the ability to capture and amplify segments containing coding sequences [11–17]. Finally, TEs can capture genes and horizontally transfer them to other species, including by the means of *Starships* in fungi or virus-like *Mavericks* in nematodes [18–23]. Exon-shuffling via the activity of TEs can lead to novel transcription factor binding sites, providing novel regulatory dynamics and ultimately new functions to proteins [24].

TE insertions into coding regions are typically deleterious and are therefore under strong purifying selection [25,26]. Yet, TE insertions into duplicated genes, pseudogenes or non-essential genes are less likely to be deleterious and may lead to neofunctionalization or exonisation of the gene [4]. If retained over time, such host-TE fusions may lose functions related to TE proliferation and become essential, a process also identified as TE domestication or co-option [24,27]. Host-TE fusions that provide essential new functions are expected to be retained, although the evolutionary timeframes of such domestication events remain poorly understood.

The initial function of TE encoding sequences is typically restricted to a few functions related to the mobilization and duplication of the elements [28,29]. Yet, how TE sequences provide additional functions for existing coding sequences remains understudied. A well-studied example of host-TE fusion is the V(D)J recombination that leads to immunoglobin diversification and provides highly conserved adaptive immunity in jawed vertebrates [30,31]. The recombination activating genes *RAG1* and *RAG2* retained mobility and can re-shuffle recombination signal sequences creating the basis for rapid sequences changes in the face of new antigens [31]. Even though the V(D)J recombination is not conserved across all vertebrates, the fusion is thought to have occurred ∼500 million years ago [31,32]. *RAG1* is a host-TE fusion gene, containing the transposase of the *Transib*-like DNA transposon and a RING finger ubiquitin ligase at the N-terminal that probably acts in dimerization and as a ligase for ubiquitination [33]. Another example is *KRABINER*, a host-TE fusion in vespertilionid bats consisting of a *Mariner* DNA transposon and *ZNF112* [34]. *KRABINER* controls the regulation of a large network of genes [34]. In the fission yeast *Schizosaccharomyces pombe*, *Abp1*, *Cbh1*, *Cbh2* are centromeric *pogo* derived host-TE fusions that led to retrotransposon silencing [35–37]. A *Bel-Pao* derived *gag* sequence was recently shown to have fused with *PEX14* gene, acquiring an intron and creating a host-TE fusion in fungi [38].

TEs are highly diverse in fungal genomes, even between closely related species, indicating independent TE activity [39–41]. TEs have played an important role in the evolution of host-associated lifestyles or local adaptation to external stress including tolerance of pesticides [42–45]. Many fungal species show distinct genome compartmentalization, featuring TE-rich and gene-poor islands and a fungal specific defense against repetitive sequences further increases the differentiation [9,46–48]. Fungi associated with animals and pathogenic lifestyles in general tend to have higher numbers of TE insertions into genes, which could either be recent insertions in non-essential genes or host-TE fusions [49]. Old TE insertions are more likely to affect genes with enzymatic rather than protein-protein interaction functions [49]. The TE content and diversity observed today may not necessarily correlate with the number of host-TE fusions, as TE activity is expected to occur in random or stress-induced bursts of proliferation [50]. Ancient and ongoing TE activity in many lineages of the fungal kingdom and the exceptional genomic resources available for such compact genomes provide a vast potential to retrace the emergence of host-TE fusions over deep evolutionary timeframes.

Here, we used a systematic approach to detect host-TE fusions in the genomes of 1,237 fungal isolates. To identify host-TE fusions, we used gene orthology and phylogenomic analyses to detect the emergence and retention of TE-derived domains in fungal proteomes. We found that TE-derived helicases are the dominant TE partner in likely host-TE fusions. The subphylum Saccharomycotina, which includes model yeasts like *Saccharomyces cerevisiae* and *Candida albicans*, shows elevated contents of host-TE fusions despite typically having small and repeat-poor genomes. Host-TE fusions are enriched for binding functions to heterocyclic compounds, organic cyclic compounds ATP, adenyl ribonuclease and adenyl nucleotide. Additionally, we identified widespread candidate host-TE fusions in ascomycetes involved in gene silencing, originating from *Helitron* and *Maverick* domains. Phylogenetic analyses suggest independent origins of identical host-TE fusions, uneven rates of gene retention and secondary losses.

## Methods

### Retrieval of genomes and gene annotations

We obtained genomes and gene annotations for 1,237 fungal isolates from two different sources. A total of 994 genomes belong to the phylum Ascomycota, 195 Basidiomycota, 28 Mucoromycota, 12 Chytridiomycota, 8 Zoopagomycota (see Supplementary Table S1 for full references and additional data). The budding yeast genomes were retrieved from Shen and colleagues [51]. We retrieved additional genomes and gene annotation from fungal and Oomycetes genomes were retrieved from NCBI. Sixteen oomycetes were used as outgroup to root the phylogenetic trees in downstream analyses (Supplementary Table S1).

### Phylogenomic reconstruction

To build a tree, we followed the approach by Li et al [52] to reconstruct the fungal tree of life. Briefly, we first identified a set of single-copy orthologous genes in each of the 1,237 genomes using BUSCO v 4.1.4 searching the fungi or oomycote orthology database version 10 for fungi and oomycetes, respectively [53]. The pipeline identified a maximum set of 756 BUSCO genes in the genome of the fungus *Colletotrichum plurivorum*. The identified BUSCO genes were then translated into protein sequences respecting the relevant genetic code (code 12 for Saccharomycotina isolates except for *Pachysolen tannophilu*s (Pactanno for which code 26 was used, and code 1 for all other genomes) [54]. Of the 756 BUSCO genes identified, a random sample of 100 of the resulting BUSCO protein sequences was then concatenated using the geneStitcher.py script (https://github.com/ballesterus/Utensils) and aligned using mafft v 7.475 with the parameters *--maxiterate 1000 --auto* [55]. The resulting alignment was then trimmed using trimAl v 1.4.rev15 with the *-gappyout* option [56]. We estimated the best-fitting evolutionary models for the concatenated 100 protein sequences using partitionfinder v 2 with the quick option *-q* and default RAxML v 8.2.12 [57,58]. The resulting partitioned model was then applied for phylogenetic inference using iqtree2 v 2.1.2 after 1,000 replicates for ultrafast bootstrap and 2 independent runs with *-B 1000 --runs 2* [59]. We rooted the tree with the non-fungal oomycete *Phytophthora parasitica* and visualized the tree using the R packages ggtree, ggtreeExtra and treeio [60–62].

### Annotation of functional domains in the proteomes

To identify putative functional domains across the analyzed proteomes, we downloaded the annotated domains hidden Markov models from the PFAM release 31 [63]. We used the hmmsearch function from the HMMER package v 3.3.2 to scan all proteomes for functional domains with the *--noali* option to speed up the process [64]. We then filtered the matching domains for a minimal bitscore of 50 and a maximal e-value of 1e-17 using the HmmPy.py script.

### Inference of trophic modes

We categorized genomes using the CATAStrophy v 0.1.0 pipeline [65]. Using the predicted proteins from all genomes, we searched for genes encoding carbohydrate-degrading enzymes (CAZymes) with dbCAN v 8 [66]. As for the PFAM annotation, we performed hmmscans on each proteome using the dbCAN hidden Markov models as query. We then applied the CATAStrophy algorithm to predict the most likely trophic mode based on the set of encoded CAZymes.

### Gene orthology analysis

We inferred gene orthology among all genomes based on protein sequence identity. We used Orthofinder v 2.4.1, which implements diamond blast v 0.9.24 for homology searches across the pool of predicted proteins [67,68]. From the initial set of 13,863,658 individual proteins encoded by all genomes combined, Orthofinder grouped 7,860,083 proteins into 299,713 orthogroups.

### Detection of candidate host-TE fusions

We retrieved previously reported PFAM domains associated with fungal TE superfamilies (Muszewska et al., 2019: see there Supplementary Table S2) and filtered for genes encoding such TE-associated PFAM domains. In a second filtering step, we removed proteins annotated exclusively with TE-associated PFAM domains. We excluded PFAM with similarity to any of the fungal TE PFAM based on SCOOP and HHSearch [69] (Supplementary Table S2). We removed all oomycete genes. We filtered out genes if the identified TE and non-TE PFAM domains had an overlap of more than 5% in the amino acid sequence. Such overlaps may indicate that the two annotations identify the same protein domain. Overlaps were identified using bedtools v 2.30.0 with the *intersect* function [70]. We retained candidate orthogroups including host-TE fusion proteins if genes encoding independent TE and non-TE PFAM domains were represented in at least five genomes and belong to the same orthogroup.

### Indication of repeat-induced point mutations

Given that host-TE fusions likely emerge after gene duplication, and gene duplication is reduced in many Ascomycete species due to repeat-induced point mutations (RIP), we compared the number of host-TE fusion candidates to the percentage of RIP affected regions of a subset of 48 genomes, previously reported [48].

### Gene ontology term enrichment analyses

We analyzed the enrichment of specific gene ontology terms among host-TE fusion genes compared to the background of all genes. To reduce the computational load, we defined the background as a 1% random subset of the entire set of genes (subset: *n* = 358,350). Gene ontology terms were assigned to genes using a GO-PFAM term translation based on Mitchell et al [71]. We created a GOAllFrame object with the AnnotationDbi package v 1.54.1 and constructed a GeneSetCollection with GSEABase v 1.54.0 [72,73]. We calculated enrichment *p*-values using the *hyperGTest* in the Category package v 2.58.0 [74]. For each MF (molecular function), BP (biological process) and CC (cellular component) Gene ontology term enrichment, a *p*-value cut-off of 1e-10 and a minimum term size of 20 was applied.

### Filtering for copy-number variation in host-TE fusion genes

To detect potential activity of the TEs represented by the identified PFAM domains in individual genomes, we analyzed potential copy-number variation of the host-TE fusion genes and their respective PFAM terms. To reduce conservatively detecting host-TE fusion genes generated by recent TE insertion events, we required host-TE fusion genes to be present in at least 5 genomes belonging to the same order and present in at least 20 genomes. Furthermore, we analyzed candidate host-TE fusion genes for their PFAM domain order along the amino acid sequence and removed orthogroups without a conserved domain order. After filtering, we extracted the predicted function of the non-TE candidate function based on the information provided in the PFAM database [63]. To remove host-TE fusion gene candidates potentially erroneously identified due to the physical proximity of genes in fungal gene clusters, we performed gene cluster analyses using antiSMASH v 3.0 using the list of predicted gene clusters from Kautsar et al [75,76].

## Results

### Phylogenetic reconstruction and genomic landscape show variable genomes in the fungal kingdom

We analyzed genomes of 1,237 fungal isolates belonging primarily to phyla of ascomycetes and basidiomycetes (Figure 1A). Based on a set of 100 single-copy genes we constructed a maximum likelihood phylogenetic tree (Figure 1B; Supplementary file F1). The tree resolves the fungal phylogeny consistently with recent analyses of similar scope [77,78]. Ascomycetes are segregated into three larger groups including the Saccharomycotina, Taphrinomycotina and Pezizomycotina. The analyzed genomes are generally of high completeness based on BUSCO analyses with a mean number of complete genes of 94.97 %, and with 95.71 % higher than 80 % (Figure 1C). The number of detected genes varied from 602 to 22,164, with generally lower gene numbers in the Saccharomycotina (Figure 1B). Assembled genome sizes were highly variable and ranged between 7.37-773.10 Mb (mean=34.43 Mb; Figure 1C). Genome-wide GC content is on average 46.4 % with an observed range between 16.3-67.8 % (Figure 1C). Genomes in Saccharomycotina, Chytridiomycota, Mucoromycota and Zoopagomycota typically exhibit GC contents below 50%.

**Figure 1:**
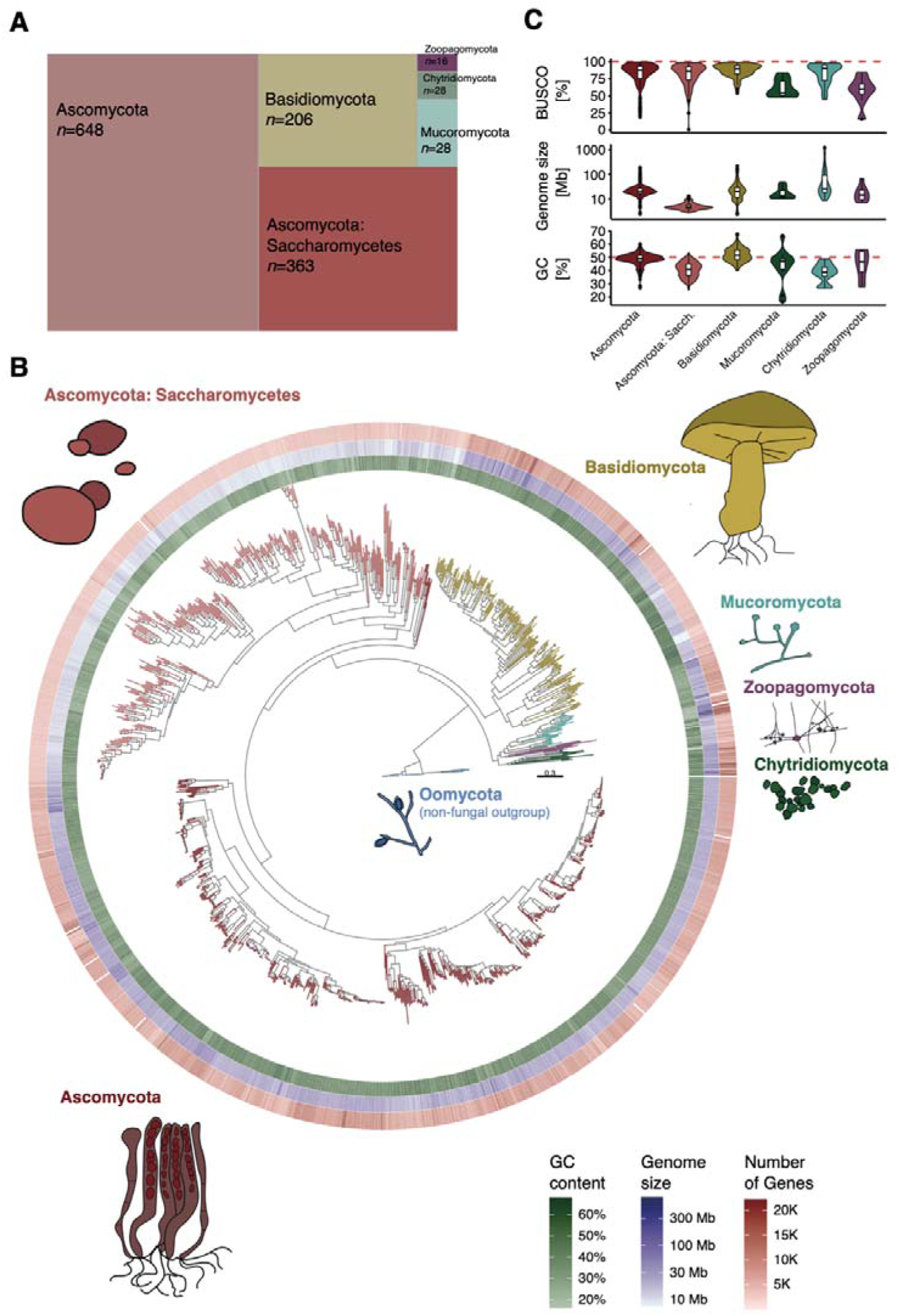
Phylogenomic analysis and genome properties across the fungal kingdom and oomycetes: (A) Number of species per phylum analyzed. The subphylum of Saccharomycotina is shown separately from the other ascomycetes. (B) Phylogenomic tree of the fungal kingdom based on 100 concatenated orthologous protein alignments. Oomycetes were defined as the outgroup. From inside to outside: genome-wide GC content (green), total genome size (blue) and number of annotated genes (red). The tree is missing the phyla of Blastocladiomycota, Cryptomycota and Microsporidia. (C) Distribution of gene completeness score (BUSCO), genome size and genome-wide GC content.

### The number of host-TE fusions across the fungal kingdom is highly variable

We next analyzed coding sequences for conserved domains present in the PFAM database. To define candidate host-TE fusion we required that at least one conserved domain matches a domain thought to be exclusively associated with TEs and at least one domain not associated with TEs. The stringent filtering, which required candidates to be detected in at least 20 genomes, allowed us to focus the analyses on conserved host-TE fusions over deep evolutionary times and to exclude pseudogenes. From a total of 39,655 unique proteins associated with a TE-associated domain across all genomes, we found 13,342 to also contain a non-TE domain (Figure 2A; Supplementary Table S3). A total of 1,205 genomes (98.3 %) carry at least one gene matching our criteria for host-TE fusion genes. We found on average 297.5 host-TE fusions (range 0-3,311) per genome (Figure 2B). Overall, 0.6-17.0 % of all annotated genes of a genome are host-TE fusions. Genomes belonging to Saccharomycotina, on average, have a higher proportion of host-TE genes per genome (240 compared to 372 across all other genomes; Figure 2B), even though they generally contain fewer genes compared to other ascomycetes (5,837 compared to 12,502; Figure 2B). Generally, Sacharomycotina have a higher percentage of genes that are containing a TE derived sequence (4.18 % compared to 3.04 %; Figure 2B). Additional outlier isolates with high proportions of host-TE fusion genes include the plant pathogens *Armillaria ostoyae* and *Microbotryum silene-dioica*, as well as the ascomycete *Fusarium poae* and the mucoromycete human pathogen *Rhizopus delemar*. We found no correlation between the number of detected host-TE fusions, BUSCO completeness scores, GC content or genome size suggesting the variation in host-TE among lineages is not meaningfully explained by variation in genome assembly quality (Supplementary Figure S1A). Repeat-induced point mutations (RIP) may impact the ability to retain duplicated sequences and early stage host-TE fusions in particular. In a subset of genomes that were previously analyzed on the strength of RIP, we detected no indication of host-TE fusions in genomes covered by more than 10% of RIP affected regions (Supplementary Figure S1B). Genomes with lower coverage of RIP affected regions vary between 0 and 10% of predicted proteins that are part of a host-TE fusion.

**Figure 2:**
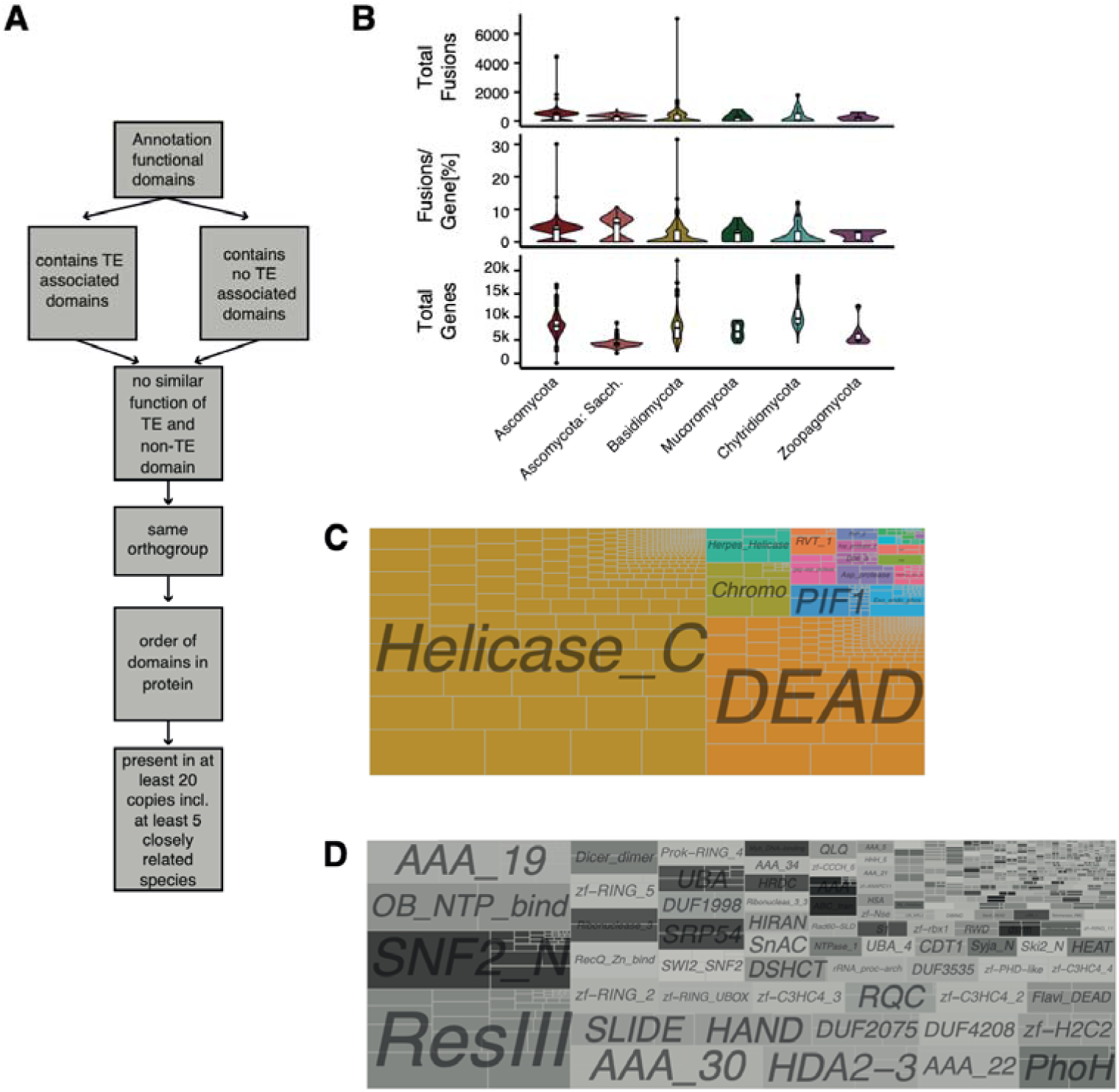
Host-TE fusion events identified across the fungal kingdom: (A) Overview of host-TE fusion detection steps. (B) Number of fusions detected per species, number of fusions detected per gene and number of annotated genes per genome. (B) Function of TE derived domains in all detected host-TE fusions. Squares indicate the number of individual fusions. (D) Function of non-TE derived domains of host-TE fusion genes.

### Transposable elements provide DNA binding sites to a wide range of functions

We restricted our analysis to 824 (115,497 occurrences) host-TE fusion orthogroups where an ortholog is present in at least five isolates, thereby retaining the evolutionarily conserved host-TE fusions. From the set of 824 individual host-TE fusions, we identified 29 distinct TE-associated PFAM (Supplementary Table S4). The TE-related PFAM only includes domains of *Helicase_C* (PF00271) *DEAD* (PF00270) helicases, *PIF1* (PF05970) and *Helitron_like_N* (PF14214) from the *AcademH*, *KolobokH* or *Helitron* TE superfamilies (Figure 2C). Non-Helicase domains with more than 1,000 candidates include *Chromo* domains (CHRromatin Organization Modifier; PF00385) from the *Maverick* TE superfamily, *Exo_endo_phos* (Endonuclease/Exonuclease/phosphatase; PF03372) from the LINE TE order and *DDE_1* (DDE superfamily endonuclease; PF03184) from *Tc1–Mariner* TE superfamily.

The diversity in non-TE PFAM domains consistently found across all orthologs of a host-TE fusion protein is substantially higher with 383 individual non-TE PFAMs. In particular, the domains included *ResIII*, *SNF2_N*, *OB_NTP_bind* and *AAA_19* functions (Figure 2D). The 383 non-TE PFAM are associated with 66 gene ontology terms with ATP binding, with highest associations in nucleosome-dependent ATP activity, nucleic acid binding, protein binding, methyltransferase activity, GTP binding, nucleus, zinc ionic binding, RNA processing and hydrolase activity, acting on acid anhydrides, in phosphorus containing anhydrides being the prevalent functions (Supplementary Figure 2A). Among our list of candidates, we find the centromere protein *CENP-B* that is known to have originated from a *pogo*-like transposase domestication event in yeast (also known as *Abp1*, *Cbh1* and *Cbh2* centromere protein N-domain in *S. pombe*; PF03184 and PF18107) [35–37,79]. More than 40 % of all non-TE domains could not be associated with a gene ontology term.

We then focused on a more restricted set of candidates including the most evolutionarily conserved host-TE fusions by requiring an ortholog to be present in at least 20 genomes (and at least five species belonging to the same order). The resulting subset of host-TE fusions contains predominantly TE-associated functions related to helicases (*Helicase_C*, *DEAD* helicase) and 241 genes encoding 125 distinct non-TE PFAM domains, including *SNF2_N*, *UBA*, *zf-H2C2*, *Rad60-SLD* and *UBA_4* (Supplementary Figure S2B). Domains with functions related to nucleotide binding are enriched in this set of 241 candidates (Figure 3; Supplementary Table S5). We also identified a fusion between a *DEAD* helicase and a *Dicer_dimerization* domain (PF00270 and PF03368). The Dicer protein is involved in RNA interference and protection against TE activity or viral infection and has been previously identified as containing a helicase domain [80].

**Figure 3:**
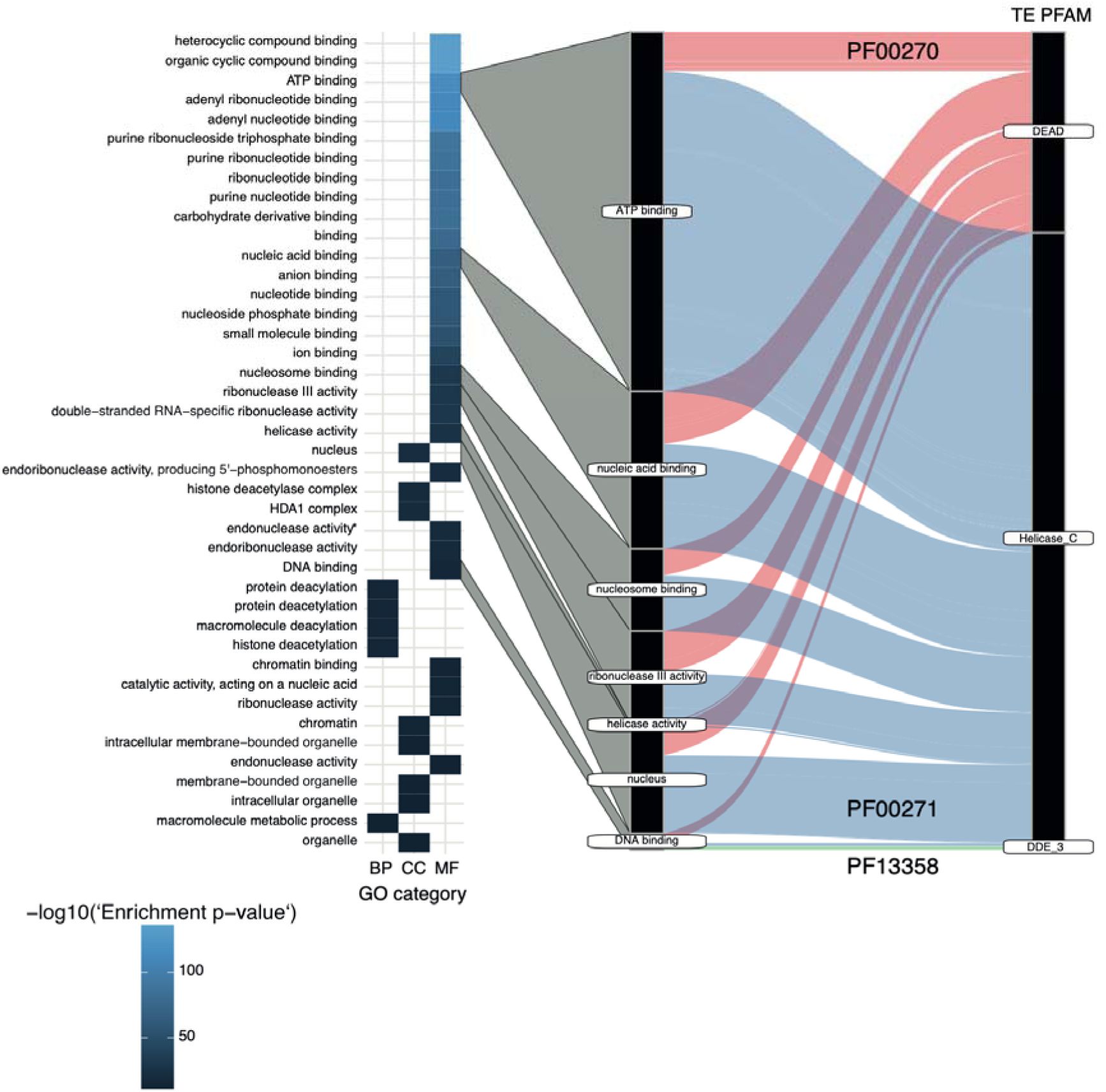
Gene enrichment analysis: gene enrichment analysis of the non-TE derived domains and the corresponding TE-derived domains. * active with either ribo- or deoxyribonucleic acids and producing 5’-p hosphomonoesters.

We highlight a host-TE fusion with homology to the Mit1 domain of an effector complex for heterochromatic transcriptional silencing (SHREC) with a function in heterochromatin silencing (PF00271 and PF00385), described in *S. pombe* [81]. SHREC is a host-TE fusion that includes a *Helicase_C* derived from *AcademH*, *KolobokH* or *Helitron*, an additional TE-derived *Chromo* domain from *Maverick* TEs and a conserved non-TE domain *zf-CCCH_6*. In addition to the conserved non-TE domain *zf-CCCH_6* and the two TE domains *Helicase_C* and *Chromo*, almost all copies of SHREC contain *SNF2_N* and *zf-PHD-like* domains. Approximately half of the fusion protein variants contain *ResII* or *PHD* domains, in addition to 88 more rarely associated domains (Figure 4A; Supplementary Figure S3). The Mit1 homology domain is primarily present in ascomycetes, with the highest representation in the Eurotiomycetes (*n*=169), Dothideomycetes (*n*=115) and Leotiomycetes (*n*=34). Lower numbers are found in Lecanoromycetes (*n*=4), Orbiliomycetes (*n*=5), Pezizomycetes (*n*=9), and Xylonomycetes (*n*=1). The Mit1 homology domain is largely absent in the large class of Saccharomycotina (*n*=1) and was only detected in *Schizosaccharomyces cryptophilus*, *S. japonicus* and *S. pombe* of the Taphrinomycotina. Weak representation is also found in basidiomycetes of the classes Agaricomycetes (*n*=4) and Dacrymycetes (*n*=1). In two ascomycetes (*Aspergillus carbonarius* and *Phialophora americana*), SHREC has a paralog, with one duplication that affected the gene with both TE domains and one duplication that affected the *Helicase_C* domain gene. A multiple sequence alignment of the duplicated genomic regions confirms the conservation of the individual domains (Figure 4B).

**Figure 4:**
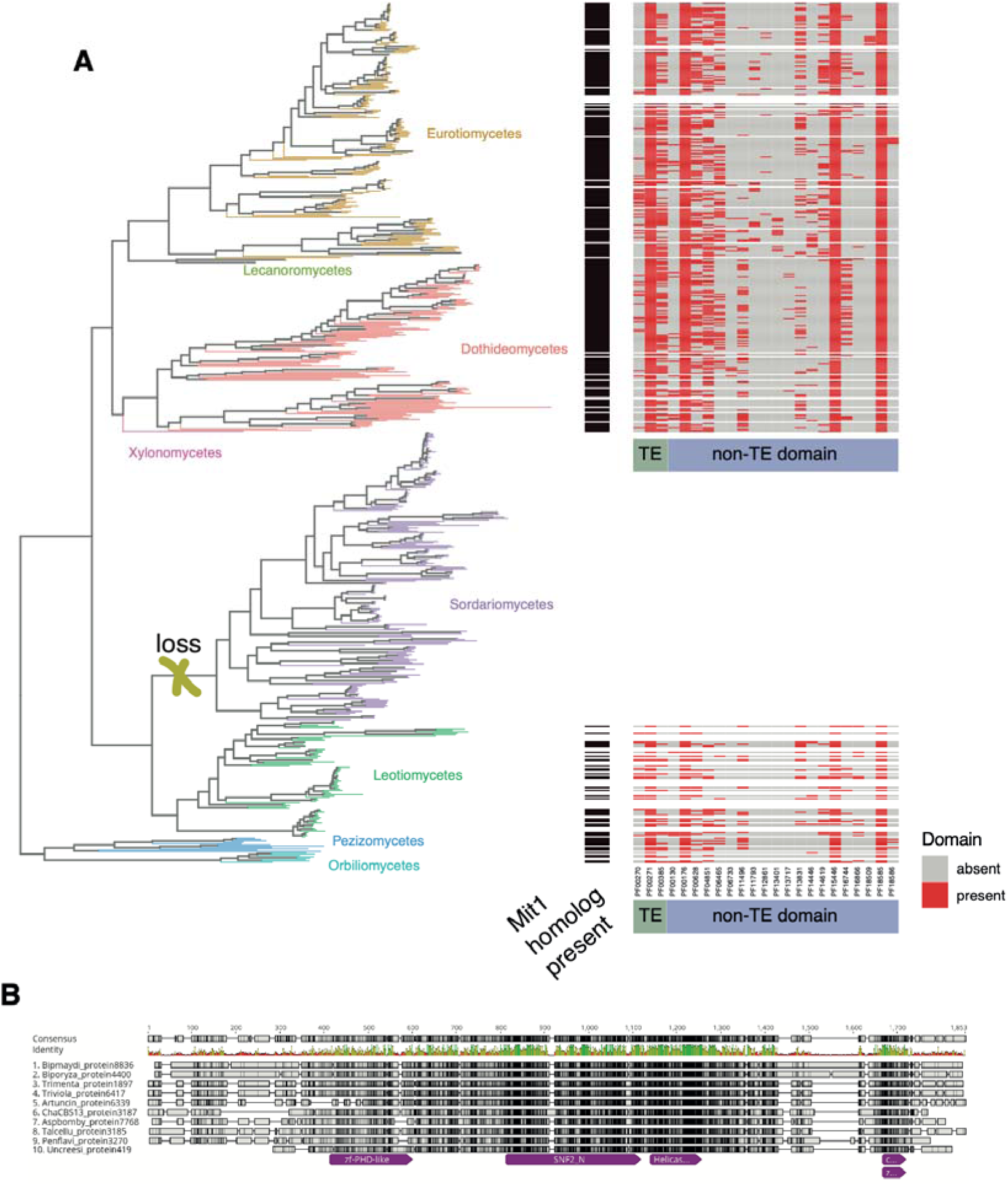
Host-TE fusion candidate Mit1 domain homolog distribution in the fungal kingdom: A) Subset of the phylogenetic tree for species with indication of a presence of Mit1 domain homologs from SHREC. The phylogenetic tree only shows Ascomycetes classes, not including the class of Saccharomycotina. Color indicates the class. Presence of the Mit1 domain homolog in the species is indicated by a black rectangle, and the presence of TE-derived domains and host-derived domains are represented with a red rectangle. B) Multiple Sequence Alignment of a selected number of proteins that are homologs to the Mit1 domain from SHREC.

## Discussion

Transposable elements are important facilitators of genome evolution, by providing regions of relaxed selection, positions of breakpoints from chromosomal rearrangements, by providing the means for horizontal gene transfer, gene mobility and reshuffling in the genome, or by providing coding regions, transcription factor binding sites and other structures to facilitate *de novo* proteins. The impact of TEs can be reversible, locally limited and short-lived. Yet, TEs may also have an impact over longer evolutionary time frames on the proteome diversification. Our analyses across 1,237 fungal genomes revealed an uneven distribution of potential host-TE fusions among major fungal phylogenies, with a higher percentage of genes involved in host-TE fusions in Saccharomycotina. Opposed to vertebrates, plants and nematodes, where terminal inverted repeat transposase domains are predominantly associated with host-TE fusions, we detected helicases as the most abundant TE-derived domains [20,27,34]. The host domains of host-TE fusions show a broader diversity in function, but tend to be associated with processes involved in genome integrity and defense against foreign sequences including TEs.

### TE-driven dynamics in the Saccharomycotina

We observed that the compact genomes of Saccharomycotina contain a higher proportion of host-TE fusions per gene compared to other fungi, while maintaining similar absolute numbers of host-TE fusions per genome. Given that all analyzed Saccharomycotina genomes have extremely low TE counts, we hypothesize that a significant proportion of detectable TEs in these species may be integrated into host-TE fusions [82–85]. Even tough TEs are rare and potentially less active than in other species, they might still play crucial roles in Saccharomycotina evolution, as seen for *CENP-B*. Notably, the absence of the ascomycete-specific defense mechanisms known as RIP (repeat-induced point mutations) against TEs in Saccharomycotina and Taphrinomycotina increases the potential for gene duplication followed by insertions of TEs and subsequent host-TE fusions events [48]. RIP is a mechanism that induces point mutations in all copies of duplicated regions of a certain length, affecting both transposable elements and genes [86–88]. RIP can introduce early stop codons or other deleterious mutations in coding regions, leading to loss-of-function of duplicated sequences [89]. Active RIP in a lineage can significantly limit the evolution of essential gene functions through gene duplication [90]. Consequently, RIP may underpin low rates of gene duplicates in ascomycetes [91]. In this context, host-TE fusions of essential genes could plausibly emerge after gene duplications, where one copy remains essential, while the other copy is under relaxed purifying selection, potentially leading to the gain of new functions through TE domain fusions. While RIP is elevating mutation rates for genes close to TEs, RIP may reduce the potential to create new host-TE fusions. With the absence of RIP in Saccharomycotina and Taphrinomycotina, host-TE fusions could arise and be retained at higher rates. Having lower numbers of host-TE fusions in genomes highly affected by RIP is an indication that this might hold true, yet the absence of RIP is not leading to high amounts of host-TE fusions.

### Helicase domains are predominant in fungal host-TE fusions

Most detected host-TE fusions encode helicase domains of likely TE origin. The most common source of helicases appears to be TEs of the DNA TE superfamilies *AcademH*, *KolobokH or Helitron* [92]. The specific *DEAD* and *Helicase_C* helicase domains were only recently recognized as of TE origin likely due to the recent discovery of *AcademH* and *KolobokH* TEs (Muszewska et al [49]. *AcademH* has been found as low-copy TEs in Basidiomycota, Ascomycota as well as Mucoromycota [92]. Helicases in general provide functions for the unwinding of DNA, DNA binding, and are involved in DNA repair pathways [93]. Helicases from *Helitrons* though are known to be able to capture neighboring regions during transposition events [12,94]. *Helitrons* might thus generate host-TE fusions through the capture of genes by a TE, rather than their own insertion into coding sequences. Once established, helicase-containing host-TE fusions might remain able to capture surrounding regions, which could be explained by a high presence-absence polymorphism of additional domains in most potential host-TE fusions involving helicases. A gene capture mechanism would also explain the high diversity of functions in host-TE fusions involving helicases. Whether the preponderance of host-TE fusions with *AcademH*, *KolobokH* or *Helitron* helicases is related to such a promiscuous mechanism to capture neighboring genes remains unknown. Recurrent gene capture by helicase containing TEs could explain the high helicase diversity in fungi and their dominance among host-TE fusion genes [94].

### A fungal host-TE fusion might be involved in silencing of repetitive regions

DNA binding activity is predominant among fungal host-TE fusion genes and is also featured among the most phylogenetically conserved fusions. The domain Mit1 in the Snf2/Hdac repressive complex (SHREC) host-TE fusion candidate shows a patchy distribution in some classes of ascomycetes, with sparse presence in other clades. Some classes of ascomycetes do not contain the Mit1 homology domain, which could be an indication that this host-TE fusion was randomly lost. In *S. pombe*, SHREC is known to transcriptionally silence genes and TEs [81]. The host-TE fusion consistently contains two TE-derived domains, *Helicase_C* and *Chromo*. *Chromo* domains are known in the superfamilies of *Maverick* (alternatively *Polinton*) and chromoviruses (a group of RLG, formerly known as *Gypsy*) and are located at the C-terminus of the integrase [95–97]. *Chromo* domains interact mostly with methylated histones [96,98]. The patchy distribution of the Mit1 homology domain and the prevalent partial loss of the *Chromo* domain indicate that the complex might not be present or not functional in all fungi, respectively [99].

Fungal proteomes have been significantly shaped by ancient and ongoing TE insertions, which may increase functional diversity and influence speciation. The exact mechanisms for creating functional proteins remain poorly documented. However, screens of populations will improve our understanding of these mechanisms. Identifying the processes responsible for creating host-TE fusions remains challenging. Regardless of genomic defenses, non-deleterious insertions of TEs into open reading frames of existing genes are likely very rare. We suggest gene capture by TEs (*i.e.*, *Helicases*) as an alternative mechanism to TE insertion into introns followed by alternative splicing to create host-TE fusion. Detecting host-TE fusions in genomes presents several challenges due to the complex nature of the events and the fact that most fusions are ancient. Accurately detecting host-TE fusions is further complicated by fragmentated genome assemblies, incomplete knowledge of TE-derived domains, and the rarity of events leading to host-TE fusions. Additionally, bioinformatics-based approaches often cannot predict novel functions of host-TE fusion genes. Future research with improved genome assembly quality, improved computational tools and functional investigations will expand our understanding or contemporary and historic host-TE fusions in the fungal kingdom.

## Acknowledgements

We thank Hadi Quesneville, Casey Bergman, Ksenia Krasileva, Anne Nakamoto and Anna Muszewska for help with TE associated PFAM. We also thank Sabina Tralamazza for help with gene cluster analysis.

## Data availability

All generated data is reported as Supplementary Tables S1-S5 and File S1.

## Author contributions

UO: Conceptualization, Data curation, Formal analysis, Investigation, Methodology, Writing; TB: Conceptualization, Data curation; DC: Conceptualization, Funding acquisition, Methodology, Project administration, Supervision, Writing

