## Supplementary Figures for "A systematic screen for co-option of transposable elements across the fungal kingdom"

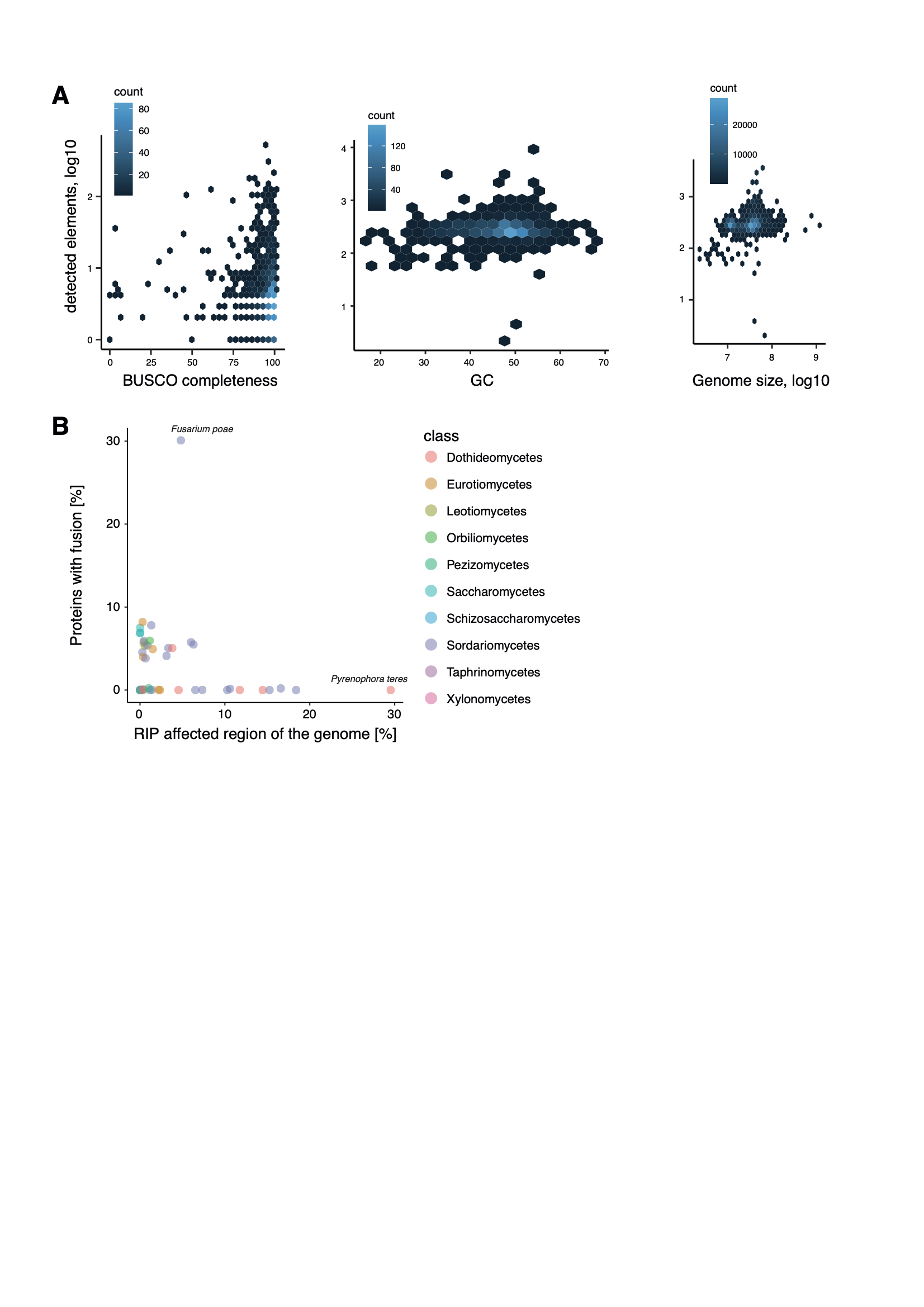

**Supplementary Figure S1: Number of host-TE detection and genome characteristics**: (A) BUSCO completeness score, genome-wide GC content, genome size. (B) RIP strength and the detection of host-TE fusions in a subset of genomes (RIP strength is the % of the genome covered by RIP affected regions; data taken from van Wyk et al,, 2021).

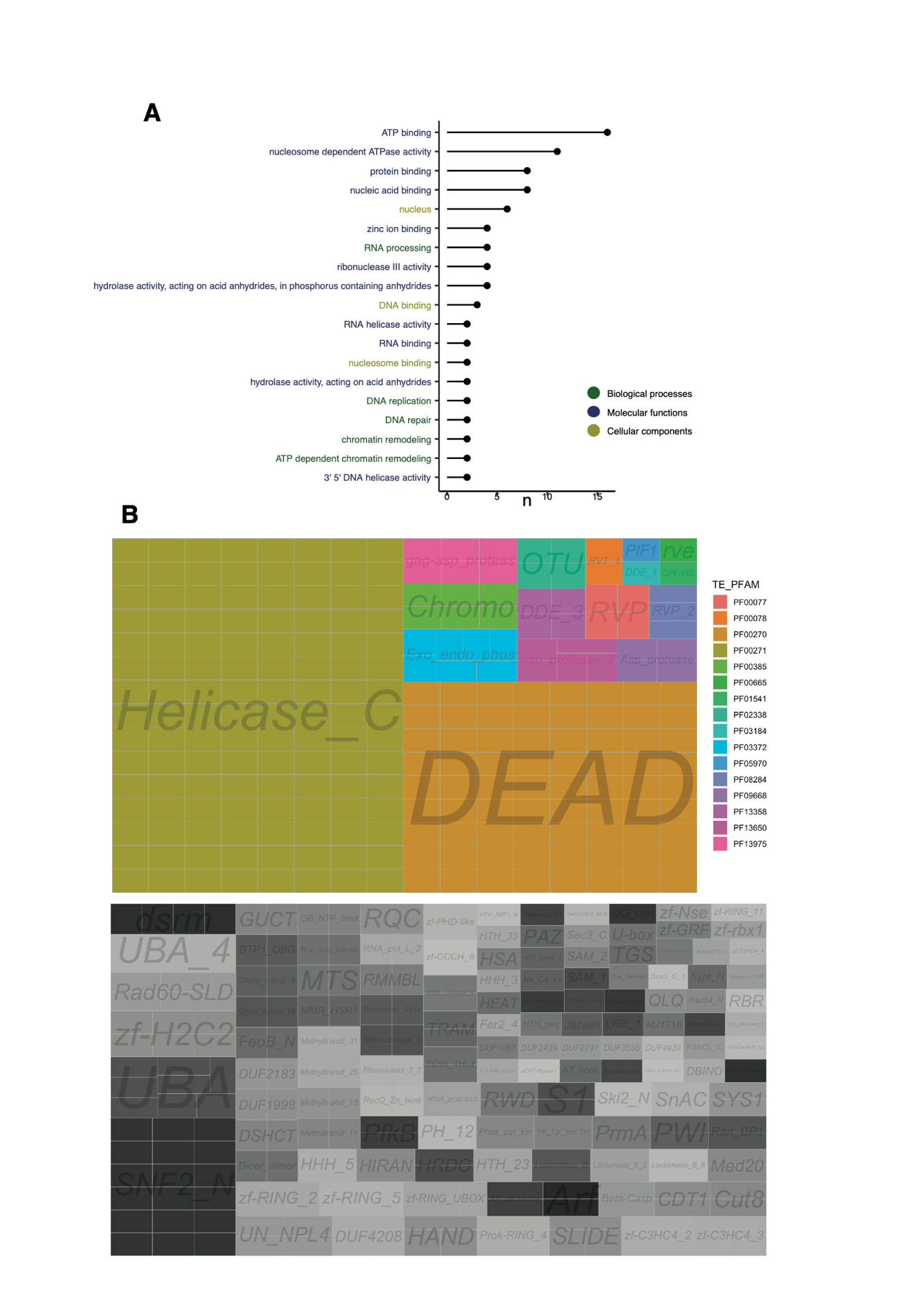

**Supplementary Figure S2: Distribution of domains in the filtered set of candidates**: (A) Counts of non-TE derived domain gene ontology terms. Gene ontology with a single occurrence can be found in Supplementary Table S5. (B) After filtering for candidates that are present in at least 20 species, with at least 5 species being closely related. Distribution for the TE-derived domain and the non-TE derived domain.

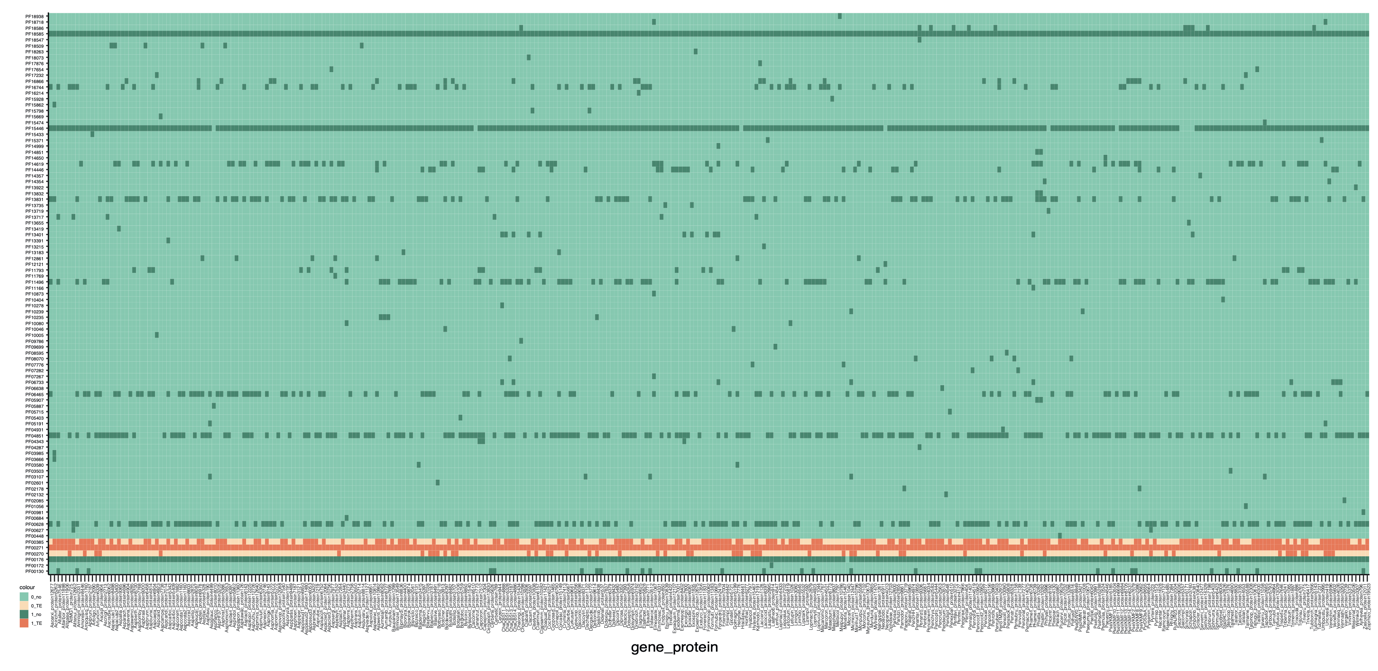

**Supplementary Figure S3: Additional domains in the host-TE fusion candidate PF00271_Can16**: Dark orange indicates presence of a TE domain in a specific gene, bright orange indicates absence of a TE domain. Dark green indicates presence of a non-TE derived domain, bright green indicates its absence.

**Supplementary File**

**Supplementary File F1: Phylogenetic tree**: phylogenetic tree of the fungal kingdom, based on 100 genes and in nexus format. The tree contains the following metadata: label (a short species ID), organism, species taxonomy ID, assembly name, assembly accession number, taxonomy ID, link to the genome in GenBank, phylum, class, order, family, genus, protein file, genome file, cds file, yeast (marked with 1 for yeast-like growing species), sequences counts, genome size (number of bases), average length, median length, maximum length, minimum length, N50, L50, BUSCO completeness score [%], BUSCO single copy genes [%], BUSCO fragmented genes [%], BUSCO missing genes [%], number of genes in the BUSCO code, BUSCO code, number of proteins detected, predicted lifestyle with CATAStrophy, phylum3 includes the differentiation between Saccharomycotina and the other classes of the phylum Ascomycota
